# Polyploidy in the adult *Drosophila* brain

**DOI:** 10.1101/852723

**Authors:** Shyama Nandakumar, Olga Grushko, Laura A. Buttitta

## Abstract

Long-lived cells such as terminally differentiated postmitotic neurons and glia must cope with the accumulation of damage over the course of an animal’s lifespan. How long-lived cells deal with ageing-related damage is poorly understood. Here we show that polyploid cells accumulate in the ageing adult fly brain and that polyploidy protects against DNA damage-induced cell death. Multiple types of neurons and glia that are diploid at eclosion, become polyploid with age in the adult *Drosophila* brain. The optic lobes exhibit the highest levels of polyploidy, associated with an elevated DNA damage response in this brain region with age. Inducing oxidative stress or exogenous DNA damage leads to an earlier onset of polyploidy, and polyploid cells in the adult brain are more resistant to DNA damage-induced cell death than diploid cells. Our results suggest polyploidy may serve a protective role for neurons and glia in ageing *Drosophila melanogaster* brains.

## Introduction

Terminally differentiated postmitotic cells such as mature neurons and glia are long-lived and must cope with the accumulation of damage over the course of an animal’s lifespan. The mechanisms used by such long-lived cells to deal with aging-related damage are poorly understood. The brain of the fruit fly *Drosophila melanogaster* is an ideal context to examine this since the fly has a relatively short lifespan and the adult fly brain is nearly entirely postmitotic with well understood development and excellent tools for genetic manipulations.

The adult central nervous system of *Drosophila melanogaster* is comprised of ∼110,000 cells, most of which are generated in the larval and early pupal stages of development from various progenitor cell types ^1,2^. By late metamorphosis, the *Drosophila* pupal brain is normally completely non-cycling and negative for markers of proliferation such as thymidine analog incorporation and mitotic markers ^13–15^

In the adult, very little neurogenesis and gliogenesis are normally observed ^12,13,16^. A population of about 40 adult neural progenitors has been reported in the optic lobe and a population of glial progenitors has been reported in the central brain ^17,18^. Upon damage or cell loss, hallmarks of cycling have been shown to be activated, although the overall level of proliferation in the adult brain remains very low ^17,18^. Thus, the brain of the adult fly is thought to be almost entirely postmitotic with most cells in G0 with a diploid (2C) DNA content. One known exception to this are the cells that constitute the “blood-brain barrier” of *Drosophila*.

The “blood-brain barrier” in *Drosophila* is made up of specialised cells called the Sub-perineurial glia (SPGs). These cells are very few in number and achieve growth without cell division by employing variant cell cycles termed endocycles, that involve DNA replication without karyokinesis or cytokinesis, as well as endomitotic cycles that involve DNA replication and karyokinesis without cytokinesis ^19,20^. The SPGs undergo these variant cell cycles to increase their size rapidly to sustain the growth of the underlying brain during larval development. The polyploidisation of these cells plays an important role in maintaining their epithelial barrier function, although it remains unclear whether these cells continue to endocycle or endomitose in the adult.

Polyploidy can also confer an increased biosynthetic capacity to cells and resistance to DNA damage induced cell death ^21–24^. Several studies have noted neurons and glia in the adult fly brain with large nuclei ^25,26^ and in some cases neurons and glia of other insect species in the adult CNS are known to be polyploid ^27^. Rare instances of neuronal polyploidy have been reported in vertebrates under normal conditions ^28^ and even in the CNS of mammals ^29,30^.

Polyploidisation is employed in response to tissue damage and helps maintain organ size ^31,32 33,34^. Therefore, polyploidy may be a strategy to deal with damage accumulated with age in the brain, a tissue with very limited cell division potential. Here we show that polyploid cells accumulate in the adult fly brain and that this proportion of polyploidy increases as the animals get older. We show that multiple types of neurons and glia which are diploid at eclosion that become polyploid specifically in the adult brain. We have found that the optic lobes of the brain contribute to most of the observed polyploidy. We also observe increased DNA damage with age, and show that inducing oxidative stress and exogenous DNA damage can lead to increased levels of polyploidy. We find that polyploid cells in the adult brain are resistant to DNA damage-induced cell death and propose a potentially protective role for polyploidy in neurons and glia in ageing *Drosophila melanogaster* brains.

## Results

Cell ploidy often scales with cell size and biosynthetic capacity ^23,35^. The brain is thought to be a notable exception to this rule, where the size of postmitotic diploid neurons and glia can be highly variable. We wondered whether alterations in ploidy during late development or early adulthood may contribute to the variability in neuronal and glial cell size in the mature *Drosophila* brain. We therefore developed a sensitive flow cytometry assay to measure DNA content in *Drosophila* pupal and adult brains. Briefly, this assay involves dissociating brains with a trypsin or collagenase based solution followed by quenching the dissociation and labeling DNA with DyeCycle Violet^36^ in the same tube, to avoid cell loss from washes. Samples are then immediately run live on a flow cytometer for analysis. We employed strict gating parameters to eliminate doublets (Supp.Fig.1A-C) ^37^. This assay is sensitive enough to measure DNA content from small subsets of cells (e.g. *Mz19-GFP* expressing neurons) from individual pupal or adult brains (Supp Fig 1D). Using this approach we confirmed that under normal culturing conditions, cell proliferation and DNA replication ceases in the pupal brain after 24h into metamorphosis (24h APF) (Supp. Fig 1E), and that only the previously described mushroom body neuroblasts continue to replicate their DNA and divide in late pupa ^14^. The brains of newly eclosed adult flies are 98-99% diploid and we, like others, only rarely observe EdU incorporation during the first week of adulthood in wild-type flies under normal conditions but not later in adulthood (Supp. Fig 1F.) ^14,16–18,38^. We were therefore surprised to find a distinct population of cells with DNA content of 4C and up to >16C appearing in brains of aged animals of various genotypes under normal culture conditions (Supp. Fig. 1G).

**Fig.1.**
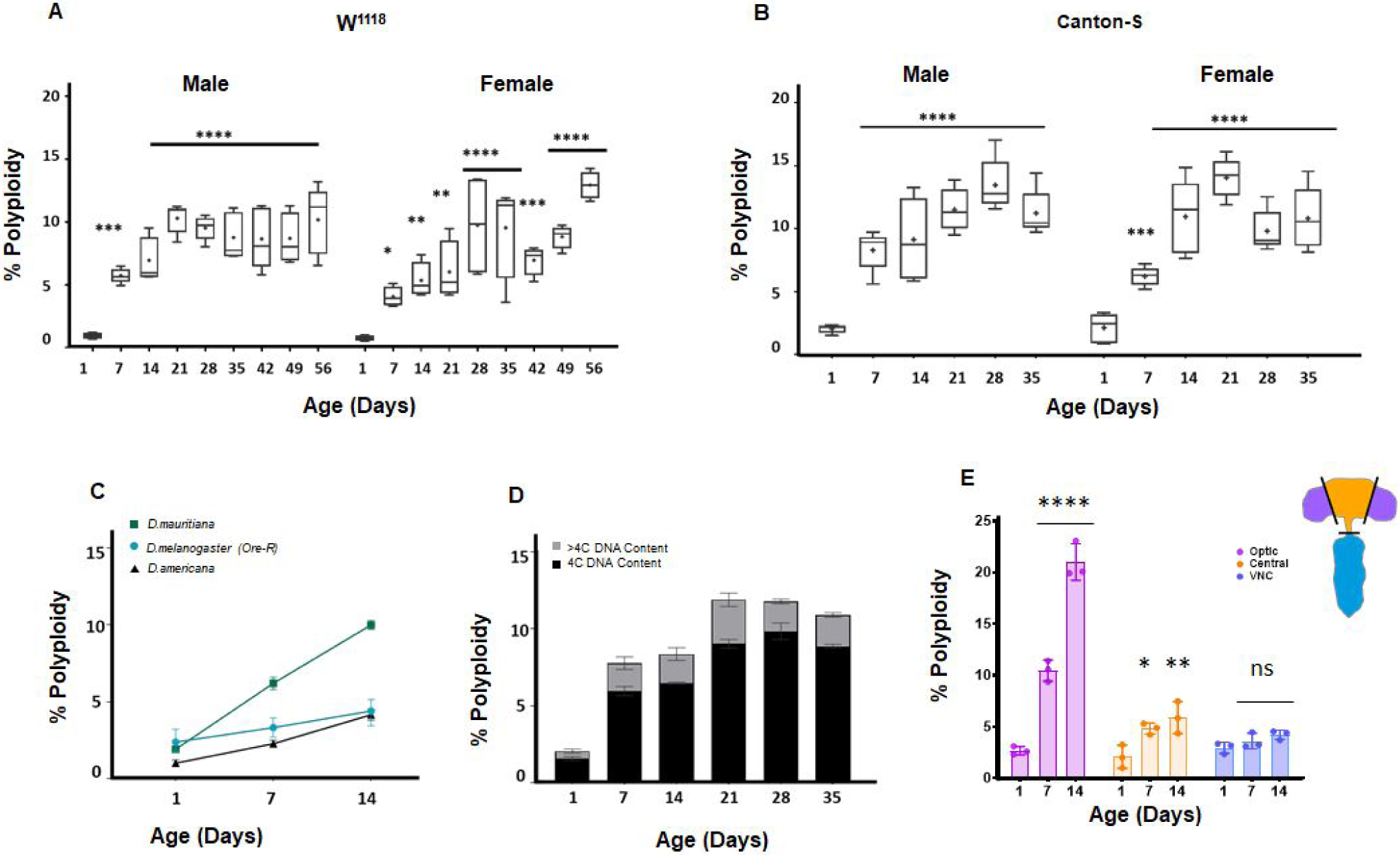
Polyploid cells accumulate in the adult *Drosophila* brain. (A,B) Percentage of cells in individual brains exhibiting polyploidy in *w*^*1118*^ (A) and Canton S (B) male and female whole brains. Age in days indicates days post-eclosion. Box plots showing range, dot indicates mean (n=10).(Two-way ANOVA with Greenhouse-Geisser correction for unequal SDs followed by Holm-Sidak’s multiple comparisons test. P values: ns>0.1234; <0.0332 *; <0.0021 **; <0.0002 ***; **** <0.0001) (C) Accumulation of polyploidy is also observed in other *Drosophila* species. *D.mauritiana and D.americana* shown respectively in green and black compared to *Oregon-R* (*D.melanogaster)* shown in teal at different time points post-eclosion. Shapes indicate mean polyploidy observed, bars show range. 3 brains each per sample, n=2 per time point. (D) Stacked bar plot showing proportion of polyploid cells with tetraploid or 4C DNA content (black) and greater than tetraploid or >4C DNA content (grey) in Canton-S males at different ages. (E) Percentage of polyploidy in (optic lobes shown in purple, central brain shown in orange and VNC shown in blue) at different ages, *w*^*1118*^ (Error bars show mean±SEM n=3).

### Polyploid cells accumulate in the adult Drosophila brain

We performed a systematic time-course to measure accumulation of polyploid cells in the adult brain in isogenic *w*^*1118*^ male and female flies cultured under standard conditions^39^. We measured the percentage of polyploid cells in individual brains from the day of eclosion until 56 days (8 weeks) at weekly intervals. Polyploid cells appear as early as 7 days into adulthood, and the proportion of polyploidy continues to rise until animals are 21 days old (Fig.1A). This increase in polyploidy is only observed until week 3, after which the proportion of polyploid cells observed remains variable from animal to animal, but, on average, does not increase (FIg.1A). We observe similar patterns of polyploidy accumulation in males and females (Fig.1A). To ensure that the polyploidy we observe is not an artefact of one particular strain, we performed similar measurements across the lifespan in other commonly used lab ‘wild-type’ strains *Canton-S* (Fig.1B) and *Oregon-R* (Fig.1C). Interestingly, *Oregon-R* flies show lower levels of polyploidy in the first two weeks than *w*^*1118*^ and *Canton-S* suggesting that different genetic backgrounds may influence polyploidy in the brain. We also performed DNA content measurements of brains from the distantly related *D.americana* which diverged ∼50 million years ago and a more closely related species, *D.mauritiana*, which diverged ∼2 million years ago (Fig.1C). While both species show accumulation of polyploidy, it is interesting to note that they show differences in levels of polyploidy.

We next measured changes in ploidy in the *D.melanogaster* adult brain over time. We pooled data from multiple animals and binned polyploid cells from *w*^*1118*^ brains into two categories: cells with ∼4C(∼tetraploid) DNA content measured by flow cytometry and cells with >4C DNA content - this includes 8C, 16C and even some 32C cells (Fig.1D). The majority of the polyploid cells appear to be tetraploid, and the fraction of cells exhibiting >4C DNA content increases during the first week of adulthood, but remains relatively consistent with age.

We next asked whether polyploid cells are located in a specific region of the brain. We dissected the *Drosophila* central nervous system (CNS) into the central brain, optic lobes, and ventral nerve cord (VNC) and measured levels of polyploidy in each region from day of eclosion to 2 weeks into adulthood (Fig 1E). We found that while there is a low level of polyploidy in the central brain and VNC that increases with age, most of the polyploidy comes from the optic lobes. Strikingly, by 3 weeks, up to 20% of the cells in the optic lobes can exhibit polyploidy.

Since the optic lobes contribute to most of the polyploidy observed, we wondered if this phenomenon may be dependent on light. *Canton-S* animals reared in complete darkness did not show difference in polyploidy compared to age-matched controls raised in regular 12-hour light / 12-hour dark cycles (Supp.Fig.2A). Next, we hypothesised that polyploidy accumulation may dependent on proper photoreceptor function. However, *glass*^*60j*^ flies devoid of photoreceptors and pigment cells in compound eyes ^40^ still show polyploidy (Supp.Fig.2B).

### Multiple cell types exhibit adult-onset polyploidy in the brain

To identify which cell types in the brain are becoming polyploid, we used the binary GAL4/UAS system to drive the expression of a nuclear-localised green or red fluorescent protein (nGRP or nRFP) with cell type-specific drivers. We then measured DNA content using dye-cycle violet in the GFP or RFP-positive populations.

We first examined the SPGs, as previous work from the Orr-Weaver lab identified these to be highly polyploid ^19,20^. When we used the SPG driver *moody-Gal4*, we found that the SPGs are highly polyploid ^19^, but contributed to less than 5% the polyploid cells observed in mature adult brains (Supp.Fig.3A,C). Another class of cells previously shown to be polyploid in some contexts are tracheal cells that carry oxygen to various organs in the fly ^41^. Using the tracheal driver *breathless-GAL4*, we found that in 10 day old adult brains, tracheal cells comprise less than 3% of all polyploid cells (Supp.Fig.3B,C). Thus, 90% of the polyploid cells we observe in the brain arise from cell types not previously known to cycle or become polyploid.

The adult fly brain is thought to be comprised almost entirely of neurons (90% of total population) and glia (10% of total population). First, we asked if neurons become polyploid by using a pan-neuronal driver, *nsyb-GAL4* to drive *UAS-nGFP* (Fig.2A). We found that indeed, by two weeks ∼5-6% of cells expressing nsyb-GAL4 show >2C DNA content. Similarly, we used the pan-glial driver *Repo-GAL4* and found that by 2 weeks ∼6-7% of glia also become polyploid in the adult brain (Fig.2B). Neurons outnumber glial cells in the fly brain, and we find that the relative proportions of polyploid cells reflect the total ratio of neurons vs. glia in the adult brain (Fig.2C). We next asked whether specific types of neurons or glia show higher levels of polyploidy. We measured the proportion of polyploid vs diploid cells in various classes of neurons (Fig.2D) and glia (Fig.2E) in 7 day old optic lobes. Interestingly, we found that most differentiated cell types we assayed in the optic lobes show some level of polyploidy by one week of age. We conclude that polyploidy arises in multiple neuronal and glial types that are initially diploid upon eclosion and become polyploid after terminal differentiation and specifically during adulthood.

**Fig.2.**
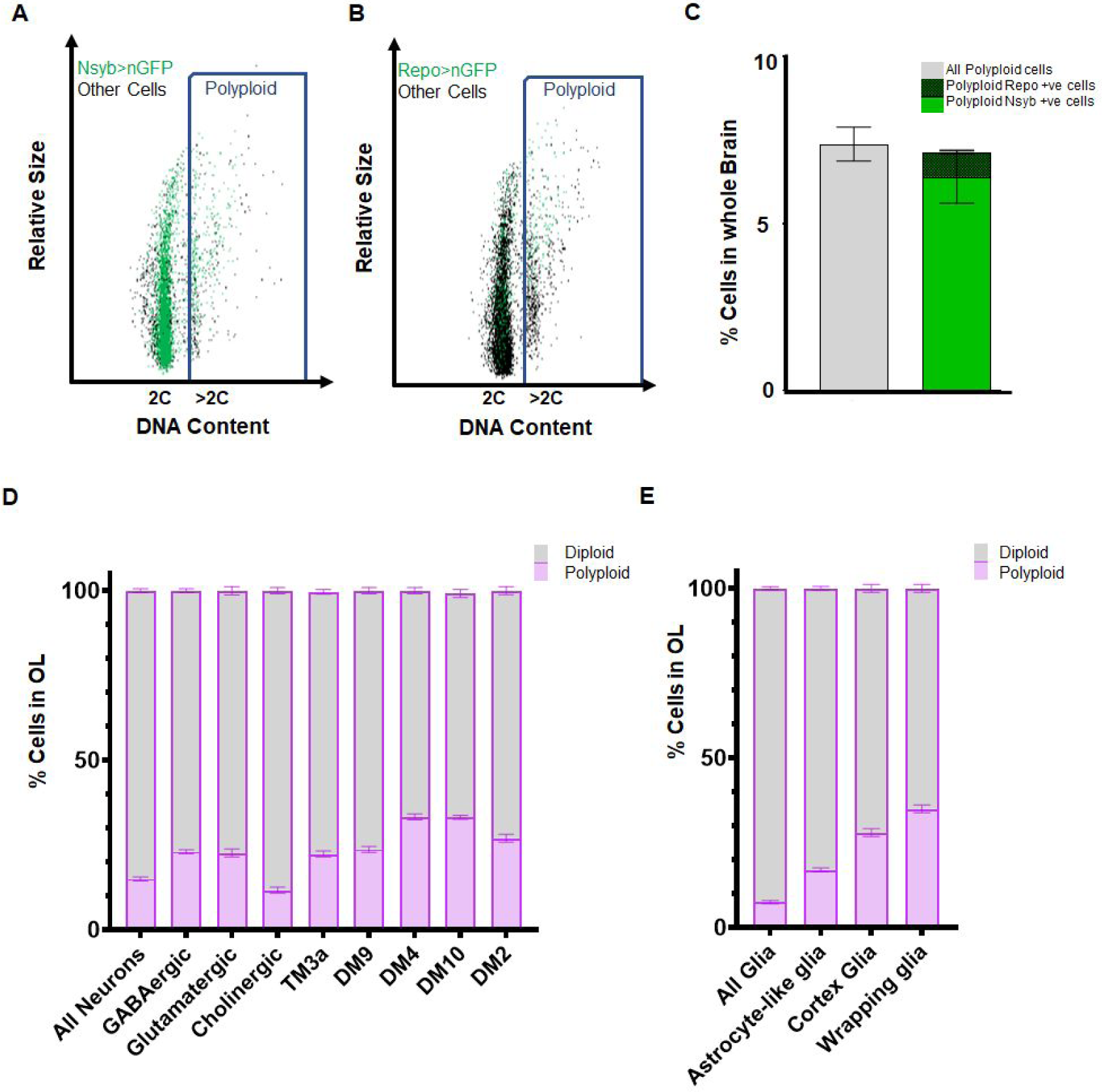
Identification of various neuronal and glial cell types that become polyploid in the adult brain. Representative flow cytometry dot plots showing polyploid neuronal (A) and glial (B) cells in 2 week old male brain(A) Neuronal nuclei are genetically labelled using nsyb-GAL4, UAS-nGFP, neuronal cells are shown in the dot plot as green dots and ‘other’ cells unlabelled by nsyb-GAL4 are shown in black. Blue rectangle highlights cells with polyploid or >2C DNA content (B) Glial nuclei are genetically labelled using Repo-GAL4, UAS-nGFP, glial cells are shown in the dot plot as green dots and ‘other’ cells unlabelled by Repo-GAL4 are shown in black. Blue rectangle highlights cells with polyploid or >2C DNA content. (C) Plot showing proportion of polyploid neurons (bold green) and polyploid glia (checked green) at 2 weeks compared to total polyploidy in the brain in W1118 control (grey) (error bars show mean ± SEM, n=3). (D) Proportion of polyploidy observed at 7 days in the optic lobes in various classes of neurons (D) and glia (E). Stacked bar plot showing mean ± SEM; percentage of polyploidy (purple) and diploidy (grey) per sample, each sample contains pooled optic lobes from 3 or more brains; (n=3)

### Polyploidy is not a result of cell fusion

We reasoned that cells in the brain could become polyploid either by re-entering the cell cycle or by undergoing cell fusion ^33,42–47^. To examine whether cell fusion occurs, we used a genetic labelling tool called COINFLP ^48^. The COINFLP genetic cassette contains two overlapping but exclusive Flippase Recombination Target (FRT) sites flanking a stop cassette that can be ‘flipped -out’ using FRT mediated recombination to give rise to cells expressing either a LexGAD driver or a GAL4 driver, which can be used to drive expression of lexA_op_-GFP (green) and UAS-RFP (red). In animals heterozygous for COINFLP, a diploid cell has only one copy of the transgenic cassette which can only be ‘flipped’ to give rise to a cell permanently labelled with either red or green fluorescent proteins, hence the name COINFLP. If labeling is induced in the brain early during development before eclosion, cells become stochastically and permanently labeled with either red or green fluorescent proteins. If cells fuse in the ageing brain, ⅓ of cells undergoing fusion could fuse a red-labeled cell with a green cell and appear yellow. We used a FLP recombinase (flippase) under the control of the *eyeless* promoter (ey-FLP) to label most of the cells red or green in the optic lobes early in development (Fig.3A-B’) and did not observe any double labelled (yellow) cells in young adult brains or older brains. We also expressed flippase enzyme more broadly under the control of a heat-shock promoter (hs-FLP) and labelled cells using a nuclear GFP or RFP at larval L2-L3 stages (Fig.3C) and measured the number of double-labelled cells in the optic lobes at on the day of eclosion or after aging at 14days post-eclosion. We never observed more than 10-12 cells per optic lobes exhibiting double-labelling under these ‘early-FLP’ conditions. These double-labelled cells under the larval hs-FLP conditions are likely the SPG cells which become polyploid early in larval development and do not express ey-FLP. We conclude that very little cell fusion occurs in the adult OL, even with age.

**Fig.3.**
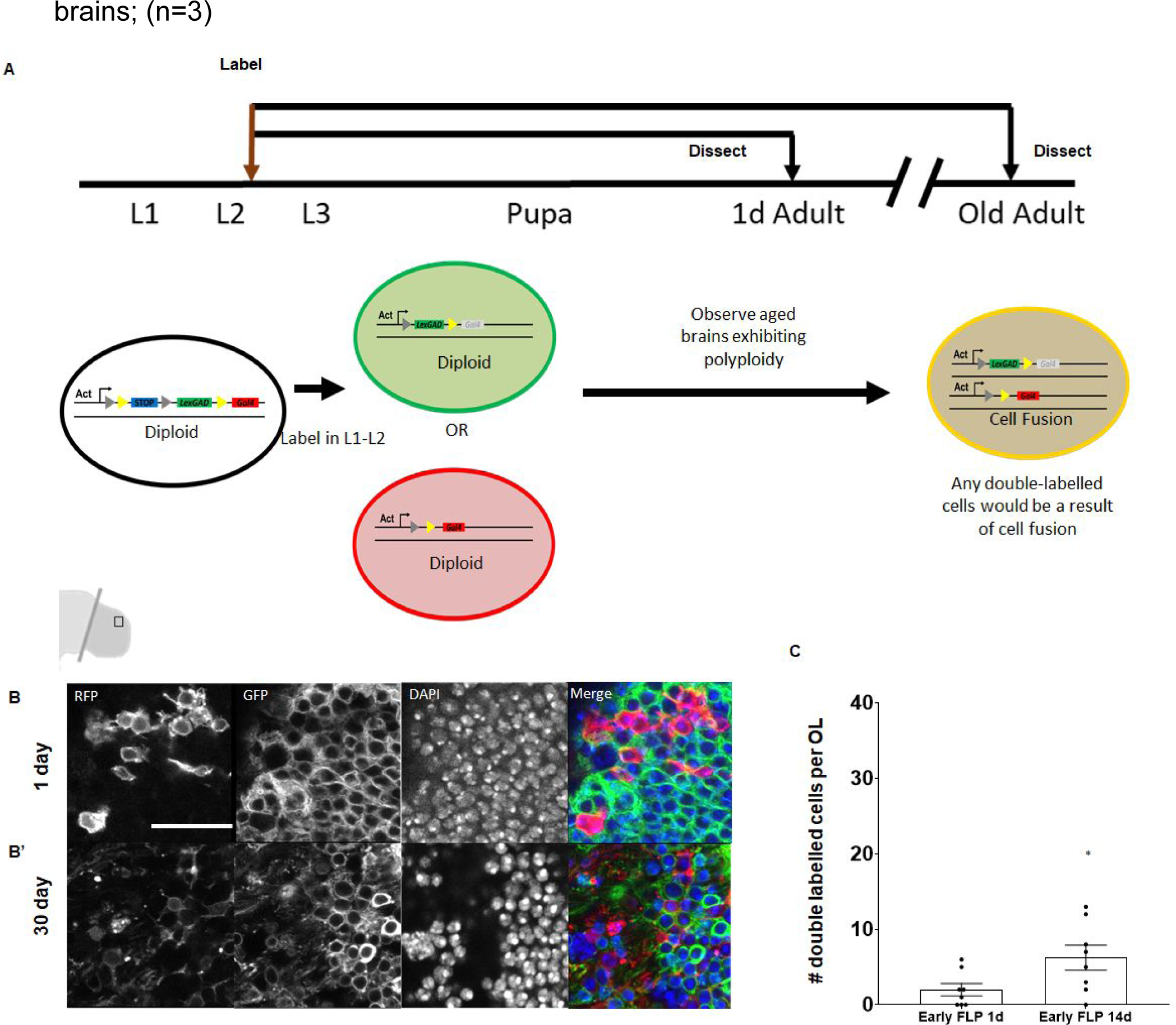
Cells in the adult brain do not undergo cell fusion to become polyploid. (A)Schematic of ‘early labeling’ using COINFLP to identify potential cell fusion events. Early COINFLP labeling will label diploid cells either with GFP or RFP. Any double-labelled cells in an older, polyploid brain will be a result of cell fusion. Representative images of 0 day (B) and 30 day (B’) polyploid optic lobes showing no double-labelled cells under ‘early-FLP’ conditions when labelled with *Ey-FLP* and membrane GFP and RFP. (C) Quantification of double labelled cells using nuclear GFP and RFP observed per brain lobe in ‘early-FLP’ condition at 14 days. P value=0.0428 significance calculated using unpaired t-test with Welch’s correction. ‘early-FLP’ in (C) was induced at L2 - early L3 stages using *hs-FLP*. Scale bars = 20µm.

### ‘Late-FLP’ can label polyploid cells that arise from cell cycle reentry

By using a modified labelling paradigm, we can also use COINFLP to label polyploid cells *in situ* (Fig.4A). Previous work with COINFLP has shown that inducing ‘flipping’ in cells that are already polyploid results in a fraction of double labelled yellow cells^48^. We therefore reasoned that heterozygous COINFLP brain cells that become polyploid by replicating their genome during ageing will contain 2 or more copies of the COINFLP transgene cassette. If we label cells by activating *hs-FLP* late in adulthood after polyploidy appears, some polyploid cells may ‘flip’ one copy green and one copy red, appearing yellow. When we induce an adult FLP at one day, before polyploidy occurs, we do not observe any double-labeled cells in the optic lobe (Fig.4B) but when we induce an adult FLP at 30 days post-eclosion, we observe several double labeled cells (Fig.4B’) indicating that these cells have undergone genome re-replication and contain multiple heterozygous copies of the COINFLP transgenic cassette. To quantify this, we used an adult ‘late-FLP’ to drive nuclear GFP and RFP, and we observe around 300 double-labelled cells per optic lobes in 14 day old brains (Fig.4C). The presence of double-labelled nuclei in aged optic lobes suggest cells become polyploid by cell cycle re-entry and endoreduplicating DNA. We can also use COINFLP labelling to identify polyploid glial cells such as cortex glia of the outer (Fig.4D) and inner (Fig.4E) optic chiasm and astrocyte-like glia (Fig.4F) in the medulla of the OL.

**Fig.4.**
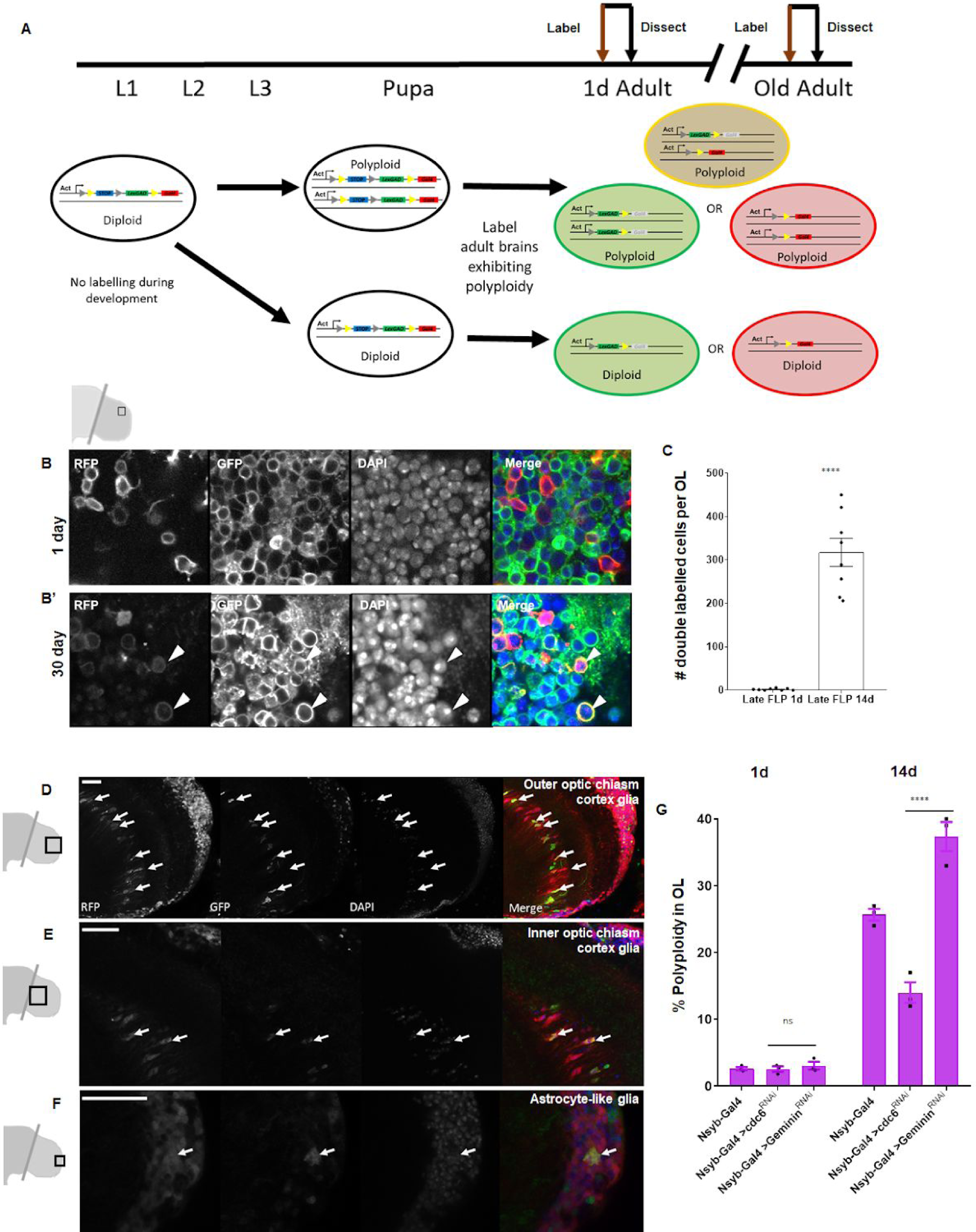
Cells in the Optic Lobes undergo cell cycle re-entry to become polyploid. (A)Schematic showing labelling protocol for inducing a ‘late flip’ in brains where polyploidy is expected to identify polyploid cells *in situ*. A proportion of cells with multiple copies of the genome will be double-labelled. Representative images of 1 day optic lobe heat shocked soon after eclosion and a 30 day old optic lobe heat shocked at 29 days to induce labelling (B). Older optic lobe shows double-labelled cells marked with membrane GFP and RFP (B’). (C) Quantification of double labelled cells using nuclear GFP and RFP observed per brain lobe in late ‘flp’ condition. Labelling was induced 24h prior to dissection for both 1d and 14d using hs-FLP. P value <.0001 significance calculated using unpaired t-test with Welch’s correction.(D-F) Representative micrographs showing cortex glia of the outer(D) and inner (E) optic chiasm as well as astrocyte-like (F) glial nuclei that can be identified as polyploid based on position and morphology using COINFLP ‘late-FLP’ labelling method. Polyploid, double-labelled glia of each type are indicated with white arrows (G) Inhibition of DNA replication licensing factor cdc6 by RNAi (BDSC55734) in neurons using the driver nsyb-GAL4 results in lower levels of polyploidy male optic lobes compared to control (GAL4 driver alone) Knockdown of replication inhibitor Geminin (BDSC30929) increases levels of polyploidy in 14 day old male optic lobes compared to 1d. Error bars show mean ± SEM, n=3. (Two way ANOVA with Greenhouse-Geisser correction for unequal SDs followed by Holm-Sidak’s multiple comparisons test p-values: 0.1234=ns; <0.0332 *; <0.0021 **; <0.0002 ***; **** <0.0001) All scale bars = 20µm.

To confirm that polyploidy in the adult optic lobes is driven by cell cycle re-entry, we used cell-type specific RNA-interference (RNAi) to modulate the DNA replication licensing factors cdc6 and Geminin in postmitotic neurons. Cdc6 is an essential factor for DNA replication licensing that promotes the recruitment of the MCM complex to load the DNA replication complex ^49^, while *geminin* is a replication licensing inhibitor that sequesters DNA replication licensing factors to inhibit DNA re-replication ^50^. Using nsyb-GAL4, we expressed *UAS-cdc6*^*RNAi*^ in differentiated neurons which significantly reduced levels of polyploidy by 14d (Fig.4G) from ∼25% in control optic lobes to ∼14% on optic lobes expressing the RNAi. We next knocked down *geminin* and found that we increase levels of polyploidy in the optic lobes (Fig.4G). This suggests that a fraction of post-mitotic neurons reactivate DNA replication to become polyploid in the ageing fly brain.

### DNA damage accumulates with age in the optic lobes

To investigate whether transcriptional changes that occur with age may be associated with cell cycle reactivation and polyploidy in the brain, we performed RNA sequencing on three parts of the brain: optic lobes, central brain and VNC from male and female *Canton-S* animals at different time points: 1 day, 2 days, 7 days and 21 days post-eclosion. To infer biological processes that are affected with age, gene ontology analysis was performed using GOrilla and redundant terms were filtered using reviGO. The most significant changes observed in the optic lobes at 21d compared to 2d are shown in Fig.5A,B. Among the most significantly upregulated groups of genes are those associated with the cell cycle, DNA damage response and DNA damage repair. The enrichment of up-regulated genes associated with the DNA damage response was also observed in the optic lobes at 7 days(Supp.Fig.4A), but the enrichment and fold-induction of specific genes is stronger at day 21(Fig.5A). A gene expression signature associated with DNA damage is specific to the optic lobes(Fig.5A, Supp.Fig.5). However the most significantly downregulated GO terms in the optic lobes(Fig.5B) at 21d are shared with the central brain (Supp.Fig.4C) and VNC (Supp.Fig.4E) and include metabolism, transmembrane transport and cellular respiration-associated processes.

To examine whether DNA damage is higher in the OL, we performed immunostaining against the phosphorylated histone 2A variant (pH2AV) in 1d optic lobes and central brain and 21d optic lobes and central brain (Fig.5C-E) from *Canton-S* male brains. Young brains show very low levels of pH2AV in both the optic lobes and central brain (Fig.5C,C’,E) but older brains show higher levels of pH2AV in the optic lobes compared to the central brain (Fig.5D,D’,E).

**Fig.5.**
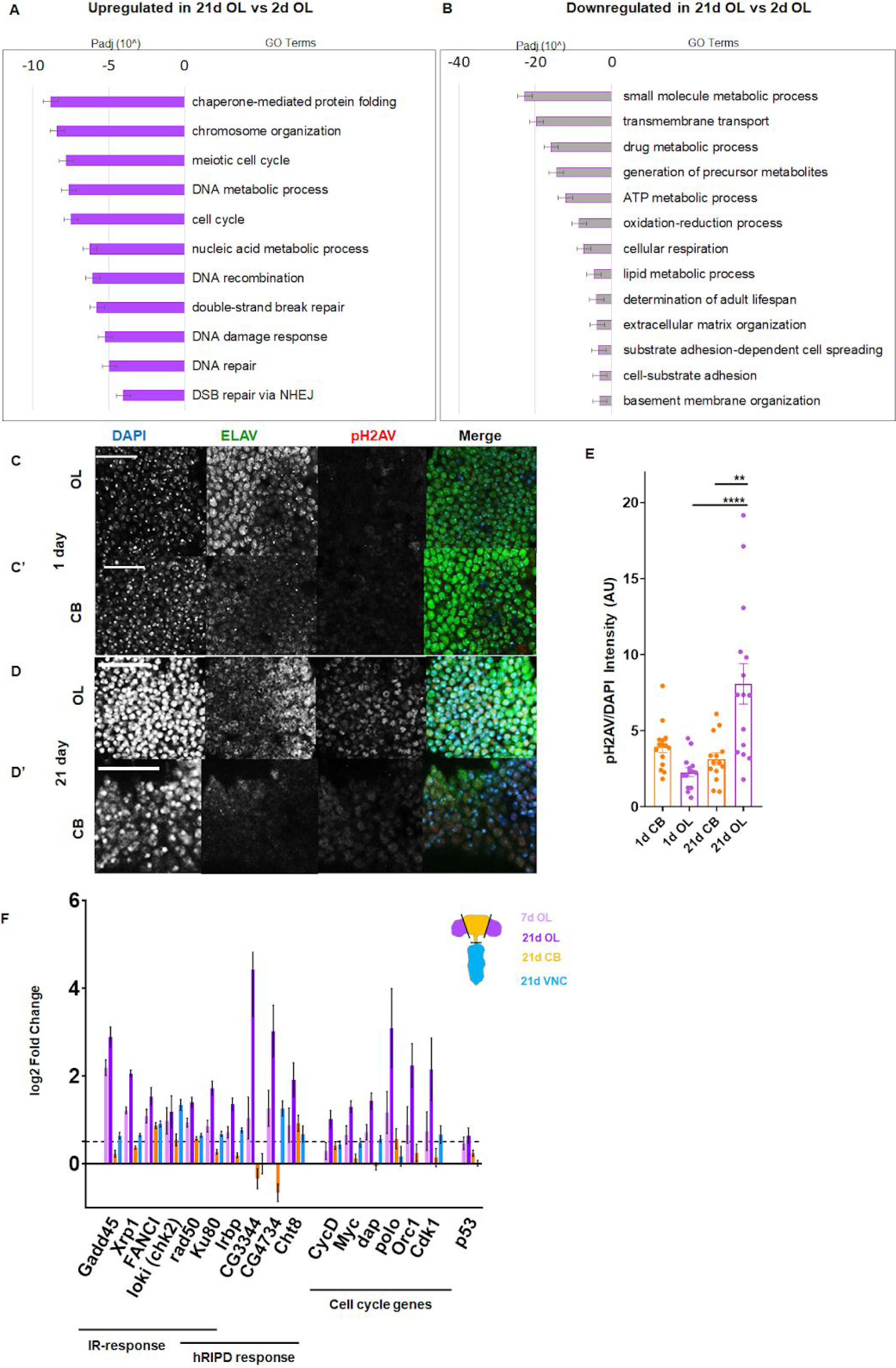
DNA damage accumulates with age in the Optic Lobes. Top changes in gene expression in 21d optic lobes compared to 2d optic lobes shown by GO term analysis. Padj= adjusted P value. Upregulated GO terms shown in solid purple (A), downregulated GO terms shown in grey bars outlined with purple (B’). Representative images showing pH2AV foci in 1 (C,C’) and 21 day (D,D’) Central Brain (CB) and Optic Lobes (OL) Neurons are labelled in green (ELAV), phosphorylated histone 2A variant (pH2AV) in red nuclei are labelled in blue (DAPI). (E) Accumulation of DNA damage is quantified by measuring pH2AV intensity/DAPI intensity per frame in 5 brains per sample. Significance determined by performing unpaired t-test with Welch’s correction for unequal SD. Scale bars = 20uM (F) Genes involved in canonical Ionising radiation (IR) response, head radiation induced p53 dependent (hRIPD) and cell cycle genes showing changes in expression compared to 2day, bars show mean ± SEM, n=3 for 7d samples and n=6 for 21d samples. Dotted line indicates threshold for significance. Genes showing changes in 7d optic lobes are shown in light purple, 21d optic lobes are shown in dark purple, 21D central brain in yellow and 21d VNC in blue.

To further understand the DNA damage and cell cycle signatures observed with age, we looked at the change in expression of specific genes involved in the DNA damage response and the cell cycle (Fig.5F). Recent work has identified a specific transcriptional response to induced DNA damage in the head that involved a non-canonical role for tumor suppressor protein p53^51^. This signature was termed head Radiation Induced p53-Dependent or hRIPD. In addition to genes such as *FANCI, loki, rad50* and *xrp1* which are involved in a canonical, ionising radiation-induced DNA damage response, we also find robust upregulation of hRIPD genes (*Ku80, Irbp, Cht8, CG3344 and CG4734*) in our RNAseq data set in older brains, specifically in 7d and 21d OL. However these genes are not as strongly upregulated in the aged central brain or VNC and *p53* itself shows only a small increase in the optic lobes at 21d (Fig.5F). Consistent with cell cycle re-entry in a fraction of cells in the OL, upregulation of cell cycle genes such as *myc, cyclin D, orc1* is observed specifically in the optic lobe sand increases with age.

### Polyploidy accumulation in neurons is p53-independent

Work in other polyploid tissues in *Drosophila* has shown that polyploid cells in the salivary gland can tolerate high levels of DNA double-strand breaks and resist apoptosis caused by DNA damage ^21,22,52,53^. This is possible because polyploid cells in tissues such as the salivary gland have intrinsically low levels of p53 protein and also suppress the expression of pro-apoptotic genes^22^. It has also been shown in various tissues and organisms that DNA damage can induce polyploidisation ^43,54,55^. Since we observe a modest upregulation of p53 as well as a p53-dependent gene expression signature in older optic lobes, we asked if the induction of polyploidy in neurons is p53 dependent. To address this, we overexpressed wildtype (p53^WT^) or a dominant-negative allele of p53 (p53^DN^) that is unable to bind to DNA and evoke a transcriptional response in neurons using the nsyb-GAL4 driver (Fig.6A). We did not see a significant difference in levels of polyploidy in 7day old brains with overexpression of either WT or mutant p53, suggesting that accumulation of polyploidy in neurons is p53-independent. We also performed cell death measurement in the brain using flow cytometry. We calculated cell death by measuring proportions of cells incorporating either Propidium Iodide (PI) or Sytox-Green (Supp.Fig.5A). We did not see a significant difference in the proportion of dead cells in overexpression of p53^WT^ or p53^DN^ conditions compared to control (Supp.Fig.5B) consistent with recent work suggesting a non-canonical, non-apoptotic role for p53 in the *Drosophila* head ^51^.

**Fig.6.**
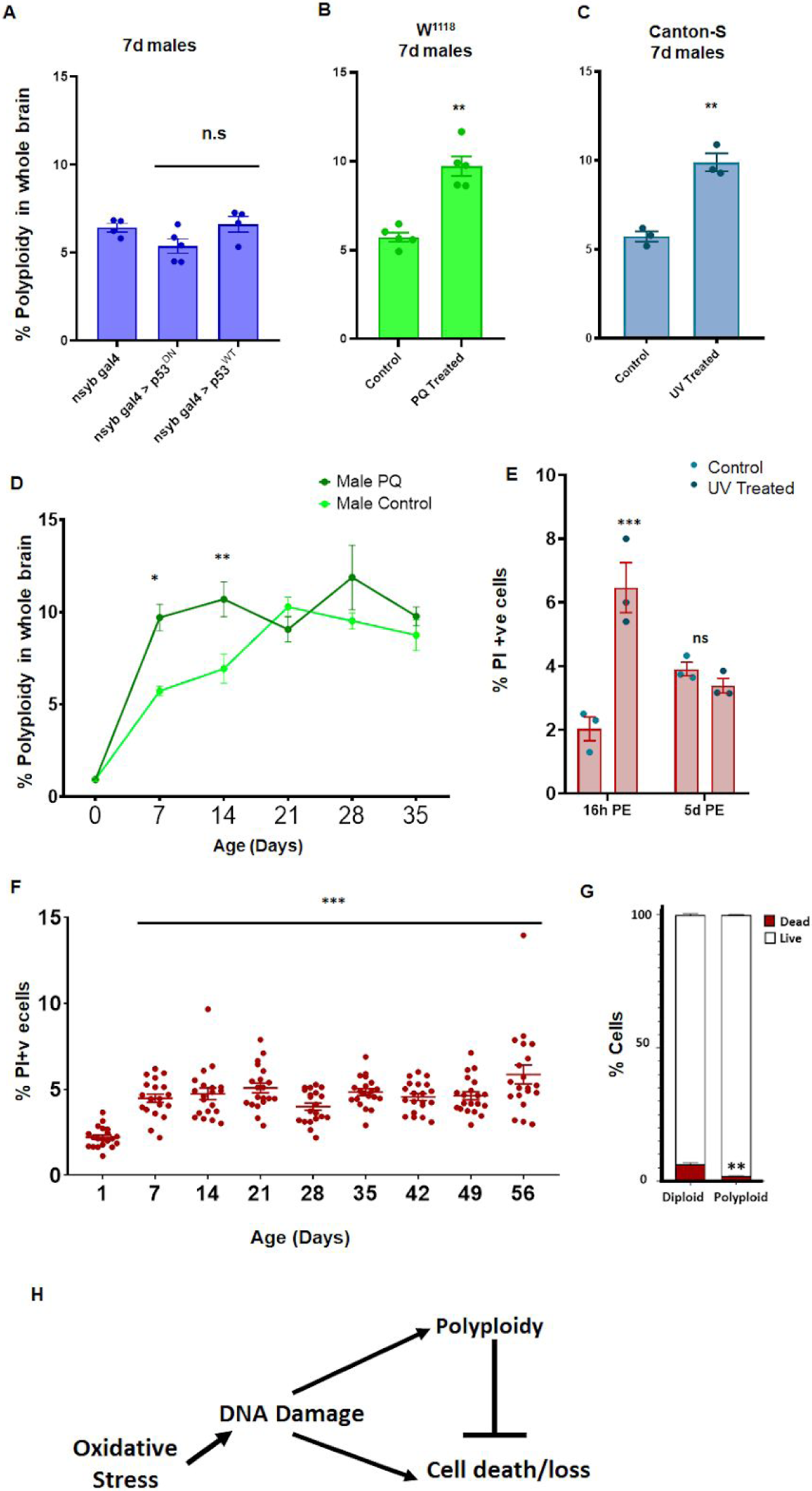
Induced oxidative stress and DNA damage results in increased polyploidy, polyploid cells are protected from cell death. (A)Higher levels of polyploidy observed after DNA damage is not P53 dependent in neurons. Percentage of polyploidy under each condition was measured in individual 7d male brains(n=5). (B) *w*^*1118*^ males treated with 2mM paraquat (PQ) from day of eclosion exhibit higher levels of polyploidy at 7 days compared to control *w*^*1118*^ males (n=5). (C) UV treated (900mJ exposure 5 days prior to dissection and flow cytometry) *Canton-S* flies show greater levels of polyploidy at 7 days compared to control (n=3). Error bars are mean±SEM, significance was calculated by performing unpaired t-test with Welch’s correction for unequal SD. (D) Accumulation of polyploidy over a time-course in *w*^*1118*^ males on 2mM PQ (dark green) compared to control *w*^*1118*^ males (light green). Shapes show mean, bars show SEM. Significance was calculated using 2 way ANOVA with Greenhouse-Geisser correction for unequal SDs, multiple comparisons with Holm-Sidak’s test; 0.1234=ns; <0.0332 *; <0.0021 **; <0.0002 ***; **** <0.000. (E) Cell death measured by Propidium Iodide incorporation in animals treated with 900mJ UV at 16h post-exposure (PE) or 5 days post-exposure. Cell death precedes accumulation of polyploidy upon induced DNA damage. Significance was calculated using 2 way ANOVA with Greenhouse-Geisser correction for unequal SDs, multiple comparisons with Holm-Sidak’s test; 0.1234=ns; <0.0332 *; <0.0021 **; <0.0002 ***; **** <0.0001 (F).PI incorporation shows percentage of dead/dying cells in individual brains, male and female, *w*^*1118*^ at different ages post-eclosion. Significance was calculated using 2 way ANOVA with Greenhouse-Geisser correction for unequal SDs, multiple comparisons with Holm-Sidak’s test; 0.1234=ns; <0.0332 *; <0.0021 **; <0.0002 ***; **** <0.0001 (G). Proportion of PI+ cells that are diploid (2C) and polyploid (>2C) in 14 day old *Canton-S* male brains n=3, Error bars are mean±SEM, significance was calculated by performing unpaired t-test with Welch’s correction for unequal SD. (H) Proposed Model

### Exogenous DNA damage leads to increased polyploidy

We next asked if exogenous stress can impact levels of polyploidy in the brain. Increased oxidative stress is commonly associated with ageing ^56–58^. We first treated flies with a low dose of paraquat (PQ) to mimic oxidative stress ^59–63^. *w*^*1118*^ flies treated with low dose of 2mM PQ from eclosion show increased DNA damage (Supp.Fig.5C) as well as increased polyploidy at 7d (Fig.6B) as well as 14d (Fig.6D) but not increased cell death (Supp.Fig.5D).

We next asked if inducing DNA damage directly affects polyploidy. We treated flies with 900mJ of UV radiation by placing flies in a UV Stratalinker at 2 days post-eclosion ^43,64^ and observed significantly increased levels of polyploidy at day 7 in UV-treated flies compared to mock-treated controls (Fig.6C). We measured cell death using propidium-iodide (PI) incorporation^36^ and observed an acute increase in cell death 16h post-exposure to UV (Fig.6E), but no difference in cell death 5days post-exposure. This suggests that cell death precedes accumulation of polyploidy upon induction of exogenous DNA damage.

### Polyploid cells are protected from cell death

Polyploid cells in other tissues are known to sustain high levels of DNA damage as well as resist cell death ^22^. We and others do not observe reproducible caspase-dependent cell death in the adult brain beyond the first 5 days after eclosion ^18^. We measured cell death and necrosis in individual *w*^*1118*^ adult brains over a time-course using PI incorporation from day 1 post-eclosion until day 56^36^. We found that newly eclosed flies exhibit a low level of dead or dying cells but from day 7 to day 56, the brain shows a relatively steady level (∼5%) of dead or dying cells (Fig.6F) although there is variability from animal to animal. Since dead cells are cleared in the brain ^65^, we expect this measurement to reflect a consistent rate of cell loss in the brain with age.

We next examined whether the polyploid cells in aged brains are protected from cell death. Since the numbers of dead or dying cells measured in individual brains was very small, we pooled brains from 2 week old *Canton-S* males to obtain a measurement of ploidy in the PI positive cells by co-staining with the DNA content dye DyeCycle Violet. We found that while ∼7% of the diploid cells incorporate PI, less than ∼2.5% of polyploid cells incorporate PI (Fig.6G), suggesting that polyploid cells are more resistant to cell death.

The work described in this study supports a model (Fig.6F) where cells in the ageing fly brain undergo endoreplication and polyploidisation in response to DNA damage and oxidative stress accumulated with age. Our data also suggests that polyploid cells are more resistant to cell death and may serve a beneficial or neuroprotective role in the ageing brain.

## Discussion

### Age-associated adult-onset polyploidy in neurons and glia

In this study we describe a surprising discovery, that diploid cells in the adult *Drosophila* brain can re-enter the cell cycle and become polyploid with age. We have identified several classes of neurons as well as glia that exhibit age-associated polyploidy. We have also characterised which regions of the brain show increased polyploidy, and find that polyploidy is closely correlated with the expression of a DNA damage response signature. Other work has also shown that a small population of stem cells in the optic lobes of *Drosophila* respond to acute injury by generating adult-born neurons ^17^. It is possible that multiple mechanisms are employed in this brain region to ensure proper function and tissue integrity with age.

Polyploidy in neurons has previously been reported in the mouse cerebral cortex ^30,66^ and chick retinal ganglion cells ^28^. Whether purkinje cells in the mammalian cerebellum are polyploid has been a matter of considerable debate over the past several decades. ^67–73^. Perhaps the most exaggerated examples of polyploidy are from the giant neurons in the terrestrial slug *Limax* ^74^ and the sea slug *Aplysia* ^75^ where giant neurons contain >100,000 copies of the diploid genome. However in all these cases, polyploid neurons appear during development. Our study describes a novel phenomenon of adult onset and accumulation of polyploidy in the *Drosophila* brain under normal physiological ageing conditions.

### What is the function of polyploidisation in the brain?

We have shown that many cell types become polyploid with in the adult brain (Fig.2). These cell types have distinct physiology and functions. How polyploidisation affects the function of these various cell types is an exciting avenue for future research. Polyploidy can confer cell-type and context specific benefits in various tissues. In *Drosophila*, polyploid intestinal enterocytes ^76^, SPGs ^19,20^ and cells in the wounded epithelium ^33,77^ undergo endoreduplication and do not undergo cytokinesis to maintain the integrity of the blood brain barrier and the cell-cell junctions in the epithelium respectively. One possibility is that polyploidy in neurons or glia may allow cells to compensate for cell loss while maintaining established cell-cell contacts. The compound eye and optic lobes of *Drosophila* contain ∼750-800 ommatidial ‘units’ that form a highly organised and crystalline structure ^5,78,79^. Numerically and topographically matched cells in the medulla cortex of the optic lobes receive inputs from the lamina which in turn receives inputs from the retina ^78,79^. We observe polyploidisation in multiple neuronal types found in the medulla, yet several cell types in the brain show a decline in number with age ^79^. In neurons, polyploidy could play a role in helping cells increase their soma size dendritic arbors ^28,80^. It is possible that polyploidy allows neurons to form more presynaptic and postsynaptic connections to compensate for lost cells while maintaining the integrity of existing connections the visual system.

Nurse cells in the egg chamber ^81,82^, cells in the accessory gland ^83,84^, salivary gland ^85^ and fat body ^86^, on the other hand become polyploid to fulfill increased biosynthetic demands. In addition to an upregulation of DNA damage and cell cycle in the optic lobes, our RNAseq data suggests compromised metabolism with age in all parts of the brain. One of the main functions of glial cells is to provide metabolic support to neurons in the brain ^87–89^. Polyploidisation in astrocyte and cortex glial cells might also serve to increase their metabolic output and compensate for the reduced metabolic output in the ageing brain.

### DNA damage accumulates in the optic lobes with age

We observe higher levels of expression of DNA damage associated genes in the optic lobes than in other parts of the brain (Fig.5A, Supp.Fig.4A). We also see higher levels of DNA damage foci in the optic lobes than the central brain. We observe this signature even at 7 days in the OL, but it becomes stronger by 21 days. This is consistent with other studies that report that signatures of ageing appear gradually over the course of an organism’s lifespan and not abruptly at later chronological ages ^90–92^. We do not know whether the increased DNA damage signature we observe in the optic lobes is because the optic lobes intrinsically sustain higher levels of DNA damage or whether other parts of the brain better equipped to resolve DNA lesions. It is also possible that we see higher levels of DNA damage foci in the optic lobes because they have more polyploid cells. Work in other tissues has shown that polyploid cells under-replicate heterochromatin and can safely harbour unresolved pH2AV foci ^21^. We see an upregulation of cell cycle-associated genes specifically in the optic lobes with age. The transcription of cell cycle genes and genes involved in the DNA damage response and repair are intimately coordinated and can be controlled by the same factors ^93,94^. Homology-directed repair of DNA lesions is only possible in S and G2 phases of the cell cycle in actively dividing cells ^95^. In other phases of the cell cycle, and after cell cycle exit, cells have to rely on error-prone non-homologous end joining mediated repair. It is tempting to speculate that re-entering the cell cycle allows postmitotic cells to repair DNA damage better and survive.

### Is polyploidy protective?

We and others observe a steady decline in number of many cell types in the adult brain with age ^18,79^. The continual loss of cells in the ageing brain may be analogous to wounding, which induces polyploidisation or compensatory cellular hypertrophy in other *Drosophila* tissues ^32,33,54 31,84^. We suggest neurons and glia in the ageing brain may employ a similar strategy, to compensate for cell loss in a non-autonomous fashion. It would be interesting to test the nature of this non-autonomous compensation by performing genetic experiments to ablate specific cell types or in sub-populations of cells.

### How does polyploidy relate to neurodegeneration?

Over the past two decades, several studies have reported an interesting correlation between neurodegeneration and cell cycle re-entry in neurons ^94,96–99^. Most of these observations are from post-mortem brains containing neurons expressing cell cycle genes or exhibiting hyperploidy (>2N DNA content). More hyperploidy is observed In brains of patients with preclinical Alzheimer’s compared to age-matched controls, which has led to the hypothesis that cell cycle re-entry may precede cell death and neurodegeneration. Whether cell cycle re-entry is a cause or a consequence of neurodegeneration has been difficult to test since both are associated with age and damage. Our data suggests that re-entry into the cell cycle may be a normal physiological response to the accumulation of DNA damage with age and that it can serve a beneficial and protective function in neurons and glia. However, we do not know how polyploidy may impact neuronal and glial function and whether it may become detrimental over time. In geriatric animals (beyond 4 weeks) we observe increased variation in the levels of polyploidy and we note that a small subset of animals also exhibit extreme levels of cell death. It is possible that these animals represent a fraction of the aged population that exhibit neurodegeneration. Our single-animal assays will be essential to identify these outliers for further study.

## Methods

### Fixation, Immunostaining and Imaging

*Drosophila* brains were dissected in 1X Phosphate buffered saline (PBS) and fixed in 4% Paraformaldehyde (PFA) in 1X PBS for 25 minutes. Tissues were permeabilised in 1X PBS+0.5% Triton-X, blocked in 1X PBS, 1% BSA 0.1% Triton-X. (PAT) Antibody staining was performed at specified concentrations in PAT (Supp.Table.1) overnight at 4°C, washed, blocked in PBT-X (1X PBS, 2% Goat serum 0.3% Triton-X) prior to incubation with secondary antibody either for 4h at RT or overnight at 4°C. DAPI staining was performed after washes, brains were wet-mounted in vectashield H1000. All imaging was performed on either a Leica SP5 or SP8 laser scanning confocal microscopes. For EdU incorporation assays, flies were placed on 10mM EdU containing food with food colouring for 3 days prior to dissection. Only flies with visibly coloured abdomens were dissected. Click-iT Plus™ staining with picolyl Azide was done as per the protocol recommended by ThermoFisher.

### Fly Husbandry

Flies were reared and aged in a protocol modified from ^39^. Ageing flies were collected soon after eclosion as virgin males and females and segregated into vials containing no more than 20 flies/vial. Ageing flies were flipped onto fresh Bloomington Cornmeal food every 5-7 days. A list of all fly stocks used in this study is supplied in Supp.Table 2.

### Image Quantification

For pH2AV quantification, 5 non-overlapping Regions of Interest (ROIs) were chosen per brain region per brain. Average Intensity of pH2AV and DAPI per ROI were computed on individual channels using ImageJ. All brains were imaged at the same laser intensity and gain settings at different ages. COINFLP double-labelled cell counting was performed manually. Individual optic lobes were imaged at 100x magnification with 100 Z-sections. Quantification was performed by cropping 5 confocal Z-sections at a time, performing maximum intensity projections of each cropped image, and counting cells that showed DAPI,GFP and RFP signal overlap.

### Heat Shock protocol

COINFLP labelling was induced with heat shock induction. Flies were placed in plastic vials and vials were then completely submerged in 37°C water bath for 15 minutes. For ‘early-FLP’, flies were moved back to room temp and dissected at day 1 or day 10. For ‘late-FLP’, heat shock induction was performed 24 hours prior to dissection. All incubations and culturing except heat shock was performed at room temperature (23°C)

### Flow cytometry

Fly brains were dissected in PBS and transferred to 1.5mL microcentrifuge tube caps containing 100uL of solution containing 9:1 Trypsin-EDTA :10XPBS with 1µL Dyecycle Violet and/or 1.12µL PI or 1µL Sytox green. Brains were incubated for 20 minutes in the microcentrifuge tube caps, triturated using low retention p200 pipette tips for 60 seconds then transferred into the microcentrifuge tubes containing 400µL of the trypsin-EDTA solution with dyes and capped, and incubated further for 45 minutes at room temperature without agitation. After incubation, each sample was diluted with 500µL 1XPBS and gently vortexed at speed 8 before being loaded onto Attune or Attune NxT flow cytometer for flow cytometry analysis.

### RNA sequencing

10 brains from *Canton-S* females and males raised at 25°C on Cornmeal/Dextrose food under normal 12h L/D cycles at ZT = 2, were dissected into optic lobes, VNC and Central brain per sample. Tissues were directly dissolved into TRIZOL-LS (Invitrogen) and RNA was prepared as directed by the manufacturer. Total RNA (2-5µg) was provided to the University of Michigan Sequencing Core for polyA selection and unstranded mRNA library preparation for the Illumina HiSeq4000 platform.

### RNAseq data analysis and GO term analysis

RNAseq analysis was performed at the U.Michigan Bioinformatics Core using the following pipeline:

1. Read files from the Sequencing Core were concatenated into single fastq files for each sample. 2. Quality of the raw reads data for each sample was checked using FastQC(version v0.11.7). 3. Adaptors and poor quality bases were trimmed from reads using bbduk from the BBTools suite (v37.90).4. Quality processed reads were aligned to the Ensembl Dm6 genome using STAR (v2.6.1a) with quantMode GeneCounts flag option set to produce gene level counts. MultiQC (v1.6a0) was run to summarize QC information for raw reads, QC processed reads, alignment, and gene count information. Differential expression analyses were carried out using DESeq2 (v1.14.1). Data were pre-filtered to remove genes with 0 counts in all samples. Normalization and differential expression was performed with DESeq2, using a negative binomial generalized linear model. Plots were generated using variations or alternative representations of native DESeq2 plotting functions, ggplot2, plotly, and other packages within the R environment.

Genes called as at least 2-fold differentially expressed between day 2 and day 21 were examined for enriched GO terms using target and background unranked lists in GOrilla and redundant GO terms were filtered using ReviGO. GO term Enrichment is presented as the -log10 of the p-value with a cutoff at p-values higher than 10^-3.

## Acknowledgements

We would like to thank members of the Buttitta & Cheng-Yu Lee labs for helpful discussions and suggestions. We also thank Yiqin Ma, Kerry Flegel, Chelsea Yu, Emily Lerner, Ashley Francis, Andrew White and Emily Rozich for help with experiments. We would also like to thank the Neuroscience and *Drosophila* research communities at UM for their support for providing fly stocks, particularly the labs of Cheng-Yu Lee, Monica Dus, Catherine Collins, Josie Clowney, Trisha Wittkopp, Scott Pletcher and Orie Shafer. This study was funded by NIH R21 AG047931, NIH R01 GM127367, ACS Scholar Award RSG-15-161-01-DDC. S.N. was supported in part by the Barbour Scholarship (University of Michigan).

---

**Sup Fig 1.**
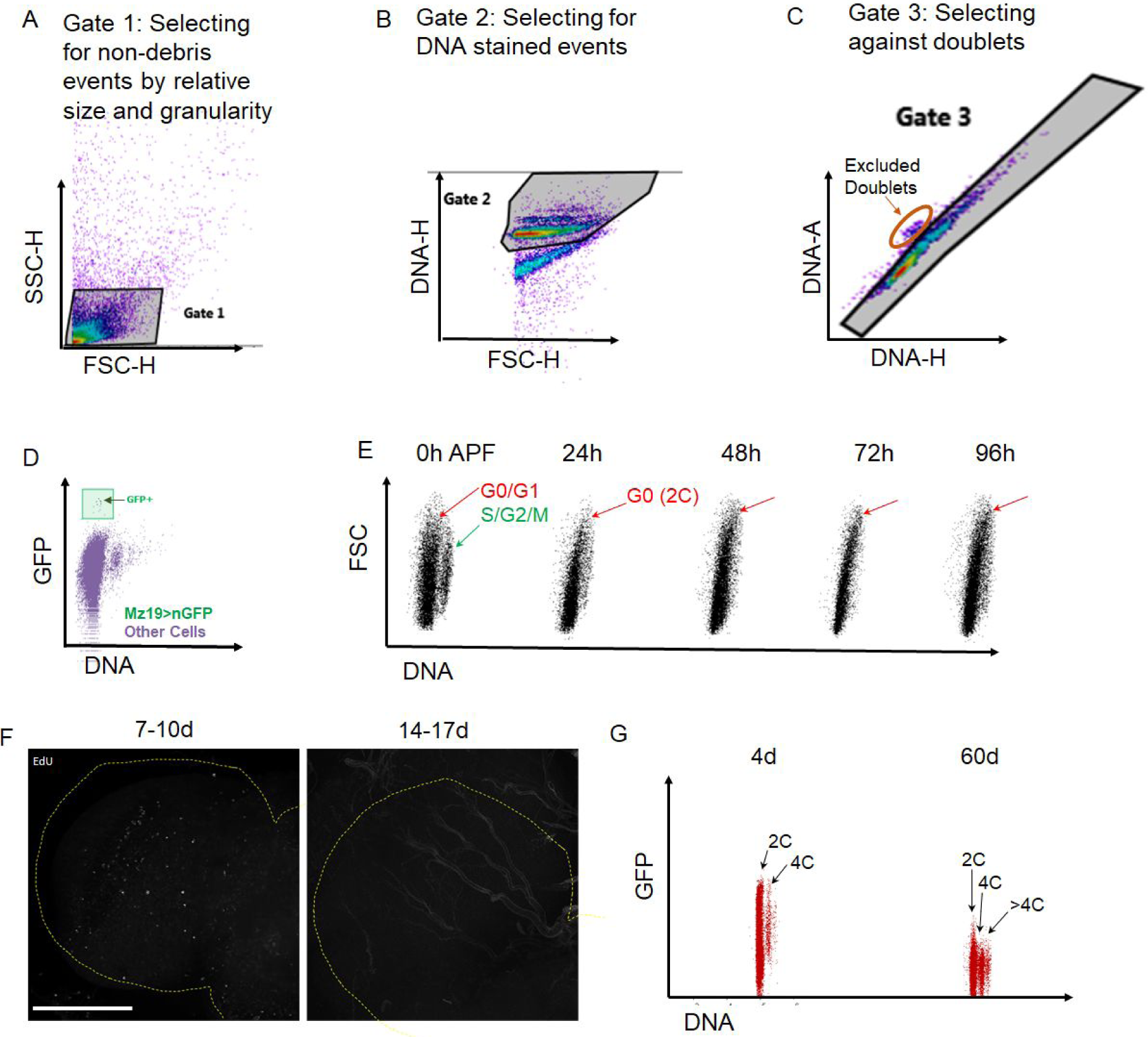
(A-C) Flow Cytometry gating strategy to ensure doublet discrimination. (D) Small population of mz19 gal4, UAS nGFP labelled neurons detectable by flow cytometry assay. (E) Dot plots showing DNA content during metamorphosis in *Drosophila melanogaster* w1118 brains. APF = after puparium formation. Cell cycle exit in most cells occurs by 24h APF. (F) Sparse EdU labelling observed in optic lobes(outlined with yellow dotted line) before 10d but not after 2 weeks in the adult brain. (G) Increased DNA content observed in 60d old brains compared to 4d. Dot plots showing all cells in Elav-Gal4, UAS-nGFP brains at each time point Polyploid cells indicated as 4C and >4C. Elav-Gal4 driver shows weak expression in older brains.

**Supp. Fig.2.**
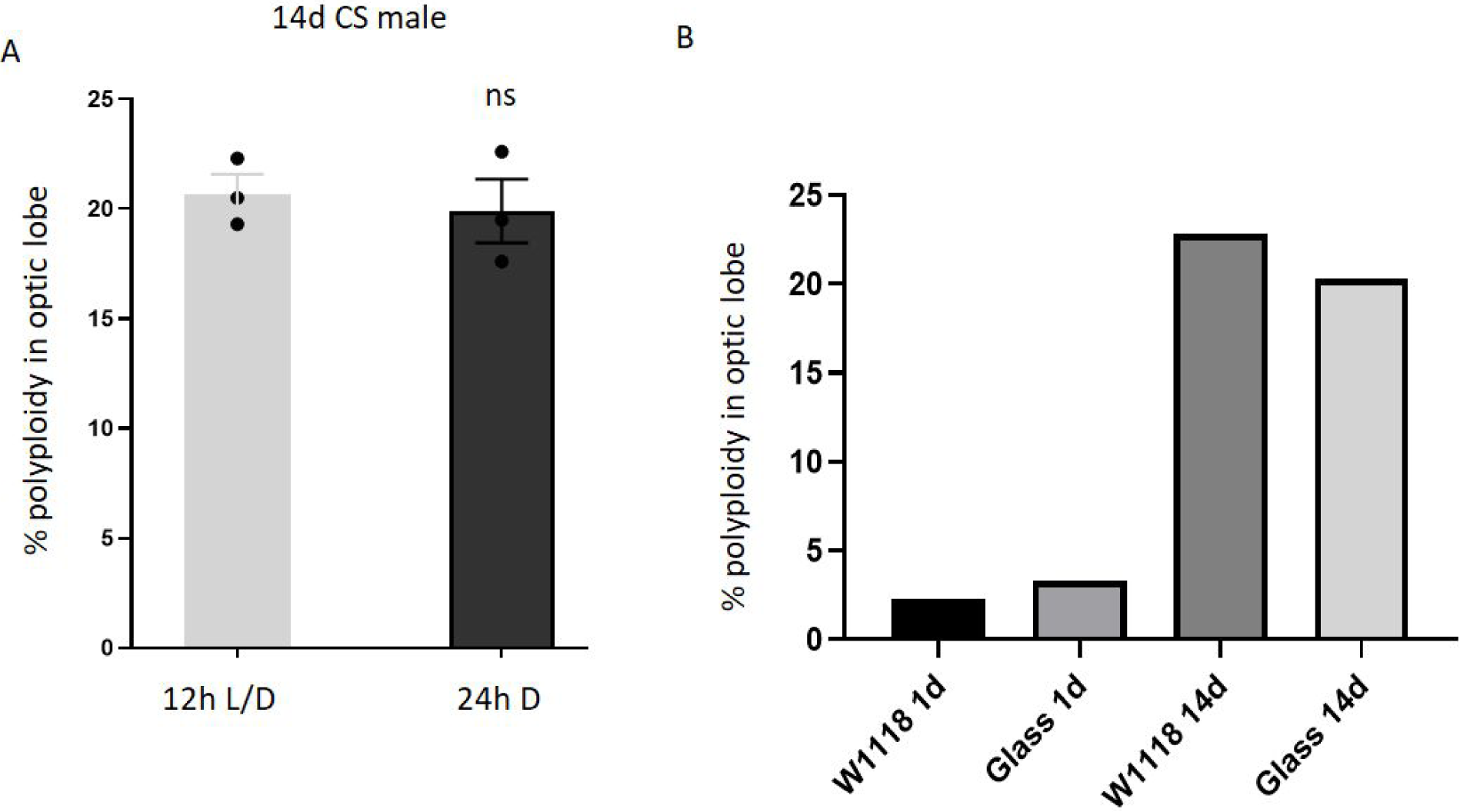
Polyploidy in Optic lobes is not light or photoreceptor dependent. (A) 14d *Canton-S* (CS) males reared in 12h L/D cycles do not show significantly different polyploidy compared to age-matched CS males reared in 24h darkness from eclosion. Error bars = mean±SEM; unpaired t-test with Welch’s correction. (B) Percentage of polyploidy observed in optic lobes of Glass 60J mutants lacking photoreceptors and age-matched *w*^*1118*^ controls at 1 and 14 days. Bars show mean % polyploidy in samples containing 3 pooled brains each.

**Sup Fig 3:**
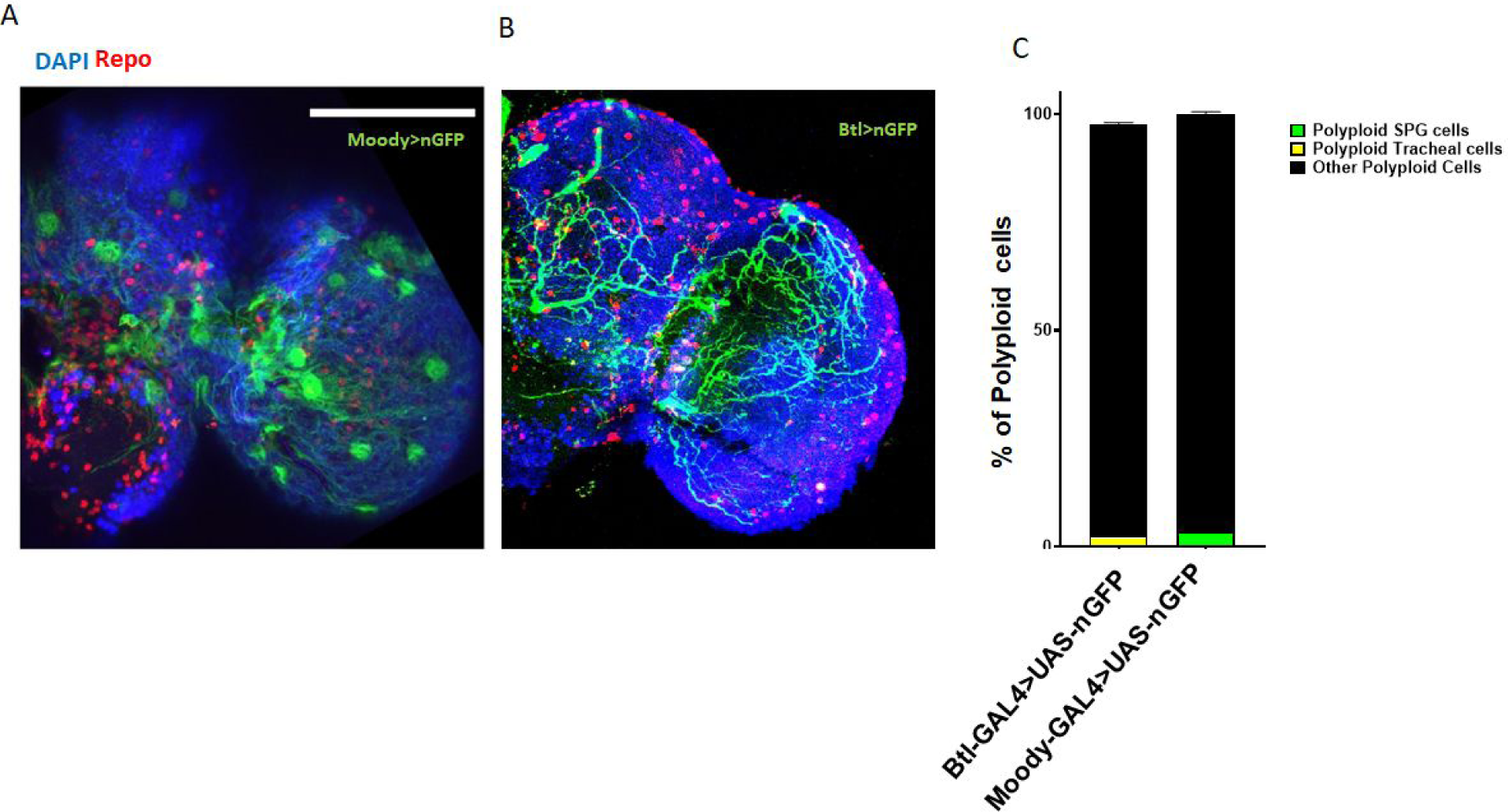
Trachea and Sub-perineurial glia comprise less than 5% of all polyploid cells. (A,B) Micrographs showing expression pattern of Moody-gal4 (A) and *Breathless Btl)-Gal4* (B) in the adult brain to label SPGs and tracheal cells respectively. (C) Flow cytometry based quantification showing the contribution of *Btl-GAL4* (yellow) and *Moody-GAL4* (green) driving cells to total polyploidy observed in 10d adult brains.

**Sup Fig 4.**
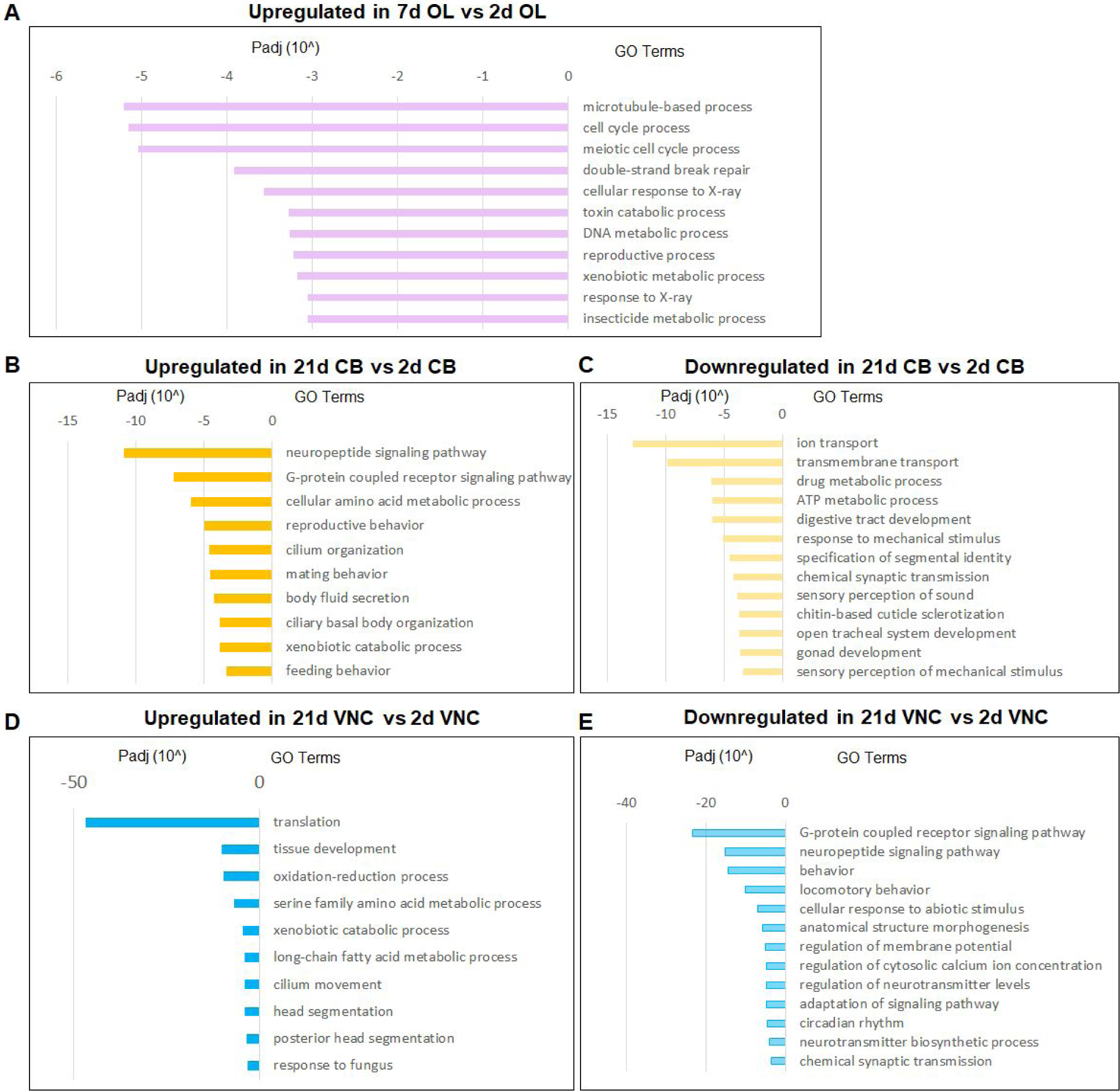
(A)Most upregulated GO terms in 7d optic lobes compared to 2d OL. Most upregulated(B) and downregulated (C) GO terms in 21d central brain compared to 2d central brain. Most upregulated(D) and downregulated (E) GO terms in 21d VNC compared to 2d VNC.

**Sup Fig 5.**
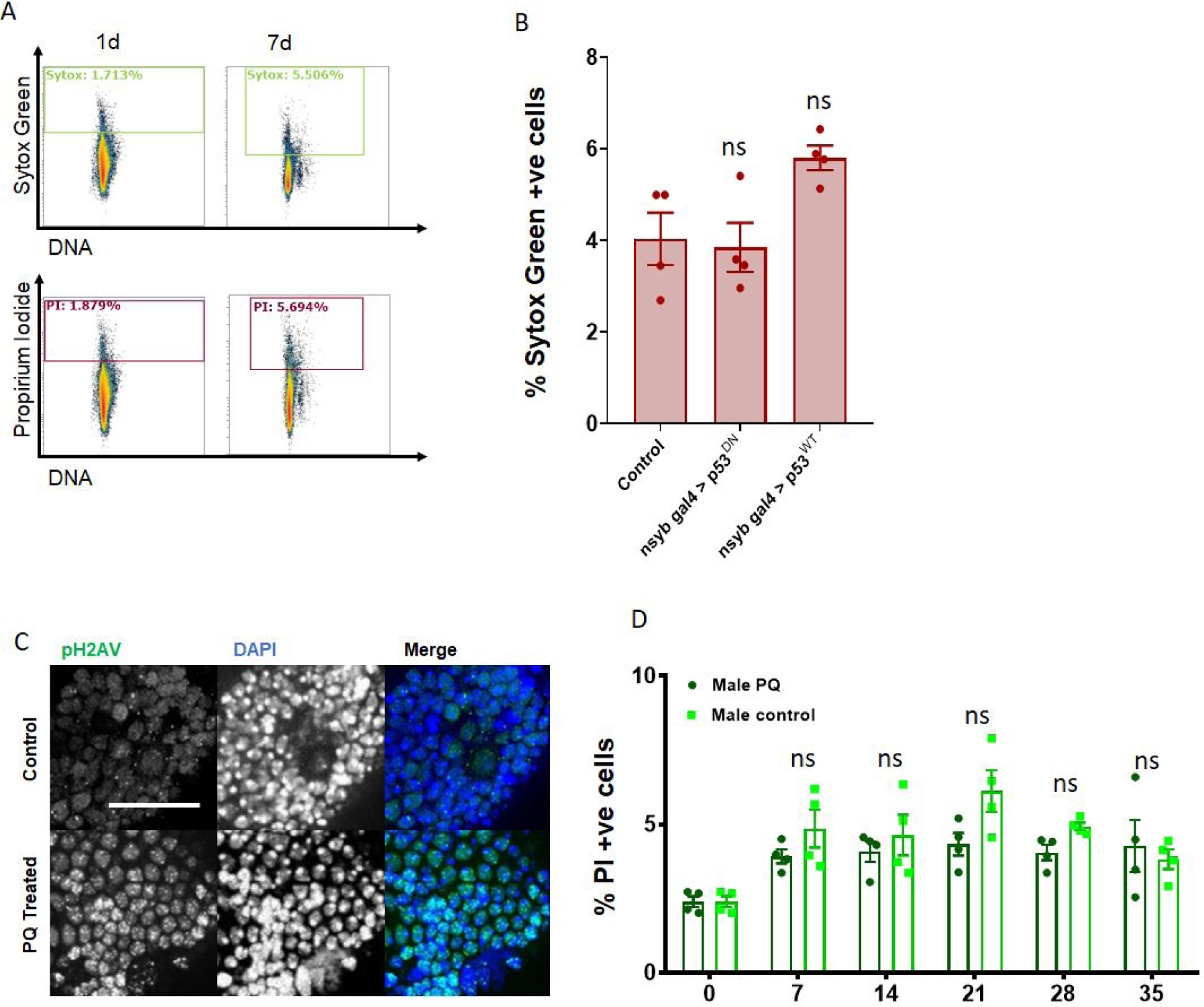
(A) 1d and 7d Canton-S brains stained with both Sytox-Green(green) and Propidium Iodide (red) show similar labelling with both cell death markers. (B) Percentage of cells incorporating Sytox-green under each condition was measured in individual 7d male brains(n=5) Error bars are mean±SEM, significance was calculated by performing unpaired t-test with Welch’s correction for unequal SD. (C) Representative images showing increased phosphorylated histone 2A variant (pH2AV) staining in 2mM PQ treated *w*^*1118*^ brain compared to age-matched control. pH2AV in green, nuclei are labelled in blue (DAPI). Scale bar=20µm (D) PI incorporation with 2mM pQ treatment measured over a time-course in *w*^*1118*^ males. 2mM PQ (dark green) compared to control *w*^*1118*^ males (light green). Error bars show SEM. Significance was calculated using 2 way ANOVA with Greenhouse-Geisser correction for unequal SDs, multiple comparisons with Holm-Sidak’s test

**Supp.Table 1.** List of antibodies and stains used.

| **Antibody/Stain** | **Concentration** | **Source** |
| --- | --- | --- |
| rat anti-ELAV | 1 : 100 | DHSB |
| mouse anti-pH2AV | 1 : 100 | DHSB |
| mouse anti-Repo | 1 : 100 | DHSB |
| DAPI | 1 : 1000 | Sigma-Aldrich |
| Alexa Fluor 568 goat anti-mouse | 1 : 1000 | ThermoFisher |
| Alexa Fluor 568 goat anti-rat | 1 : 1000 | ThermoFisher |
| Alexa Fluor 488 goat anti-mouse | 1 : 1000 | ThermoFisher |
| Alexa Fluor 488 goat anti-rat | 1 : 1000 | ThermoFisher |
| Dye-cycle violet | 2μL in 1000μL | ThermoFisher |
| Sytox Green | 2μL in 1000μL | ThermoFisher |
| Propidium Iodide | 2.25μL in 1000μL | Sigma-Aldrich |

**Supp.Table 2.**
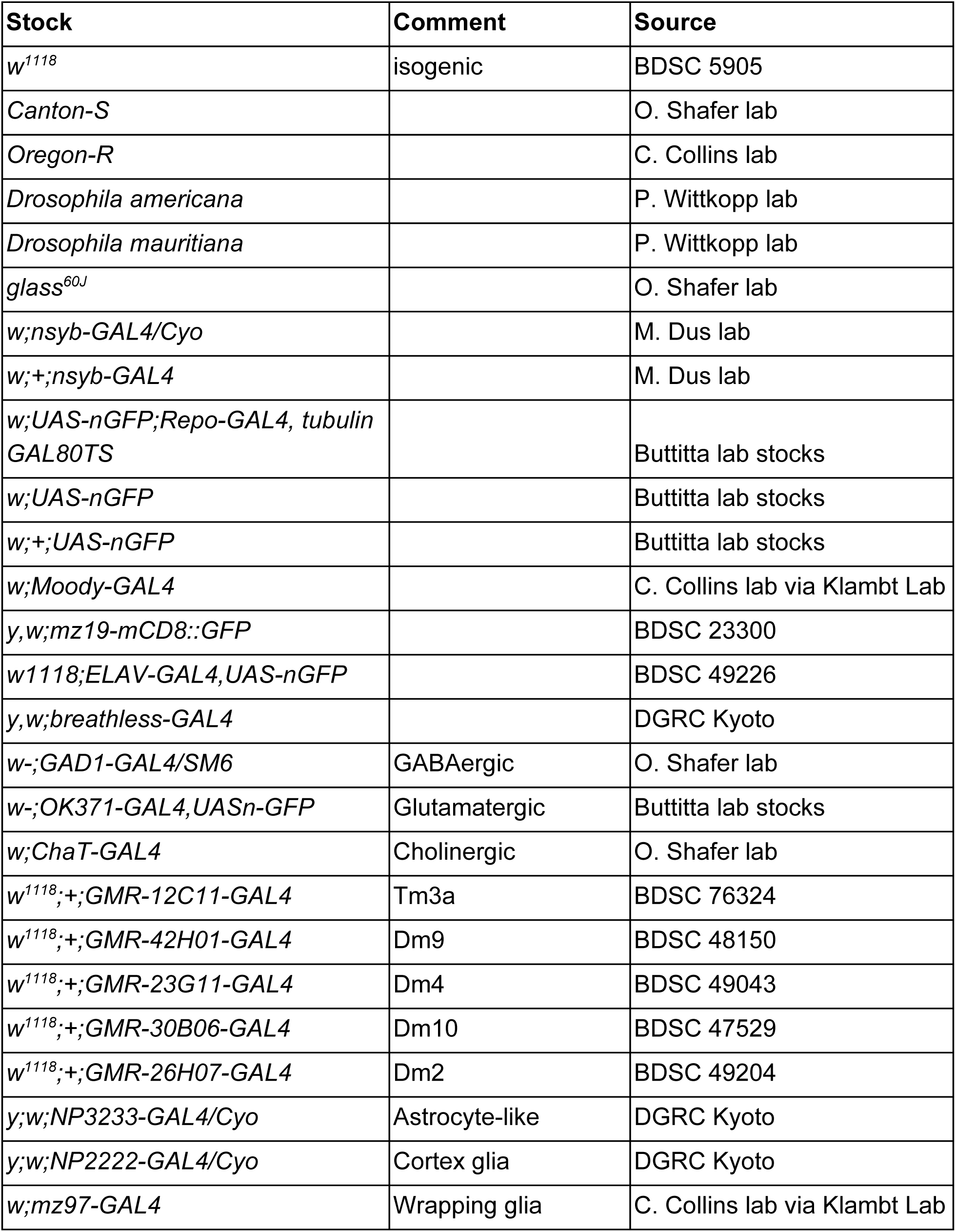
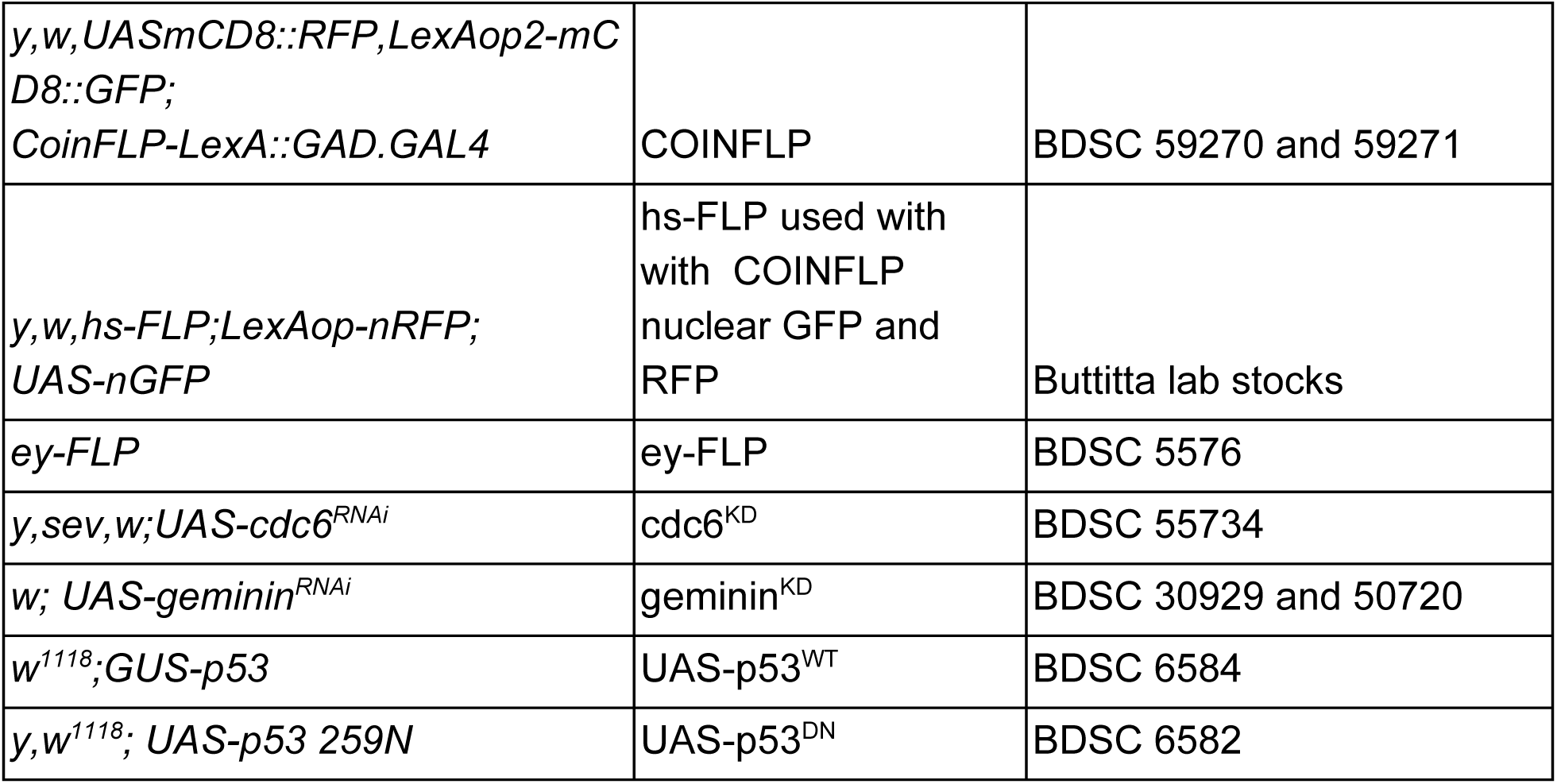
List of Fly Stocks.

